# Crystallizing the Parkinson’s Disease Protein LRRK2 Under Microgravity Conditions

**DOI:** 10.1101/259655

**Authors:** Sebastian Mathea, Marco Baptista, Paul Reichert, April Spinale, Jian Wu, Marc Allaire, Brian Fiske, Stefan Knapp

## Abstract

Mutations in the gene coding for leucine-rich repeat kinase 2 (LRRK2) are a considerable cause for Parkinson’s disease (PD). However, the high- resolution 3D structure of the protein is still lacking. This structure will not only help to understand PD etiology but will also enable rational drug design. We have established a reliable method to produce LRRK2 crystals for the first time. However, the limited resolution of the diffraction data prevented structure determination using crystallographic methods. Herein we describe our efforts to improve the crystal quality by crystallizing under microgravity conditions aboard the International Space Station (ISS). Our method features diffusive sample mixing in capillaries and controlled crystal formation by transporting the samples in a frozen state. The crystallisation was successfully repeated under microgravity conditions. However, comparison of earth-grown and microgravity-grown LRRK2 crystals did not reveal any differences in diffraction quality. Here we present the established protocol and our experience adapting crystallization condition to the requirements necessary for successful crystallization of large and sensitive biomolecules under microgravity.

## Introduction

### LRRK2 in cell physiology

The human leucine-rich repeat kinase 2 (LRRK2) is a unique protein, both in its modular architecture and in how it functions in the cell. LRRK2 comprises seven distinct domains (**Figure 1A**), most of which are thought to mediate scaffolding, protein interactions and subcellular localisation. The other two domains, namely the GTPase and the kinase domain, possess catalytic activity. While the LRRK2 GTPase acts as a conformational switch^*1*^, the LRRK2 kinase phosphorylates Ser/Thr residues of its cellular substrates. These were recently identified to be Rab GTPases^*2*^, which in turn modulate all aspects of membrane trafficking^*3*^.

**Figure 1.**
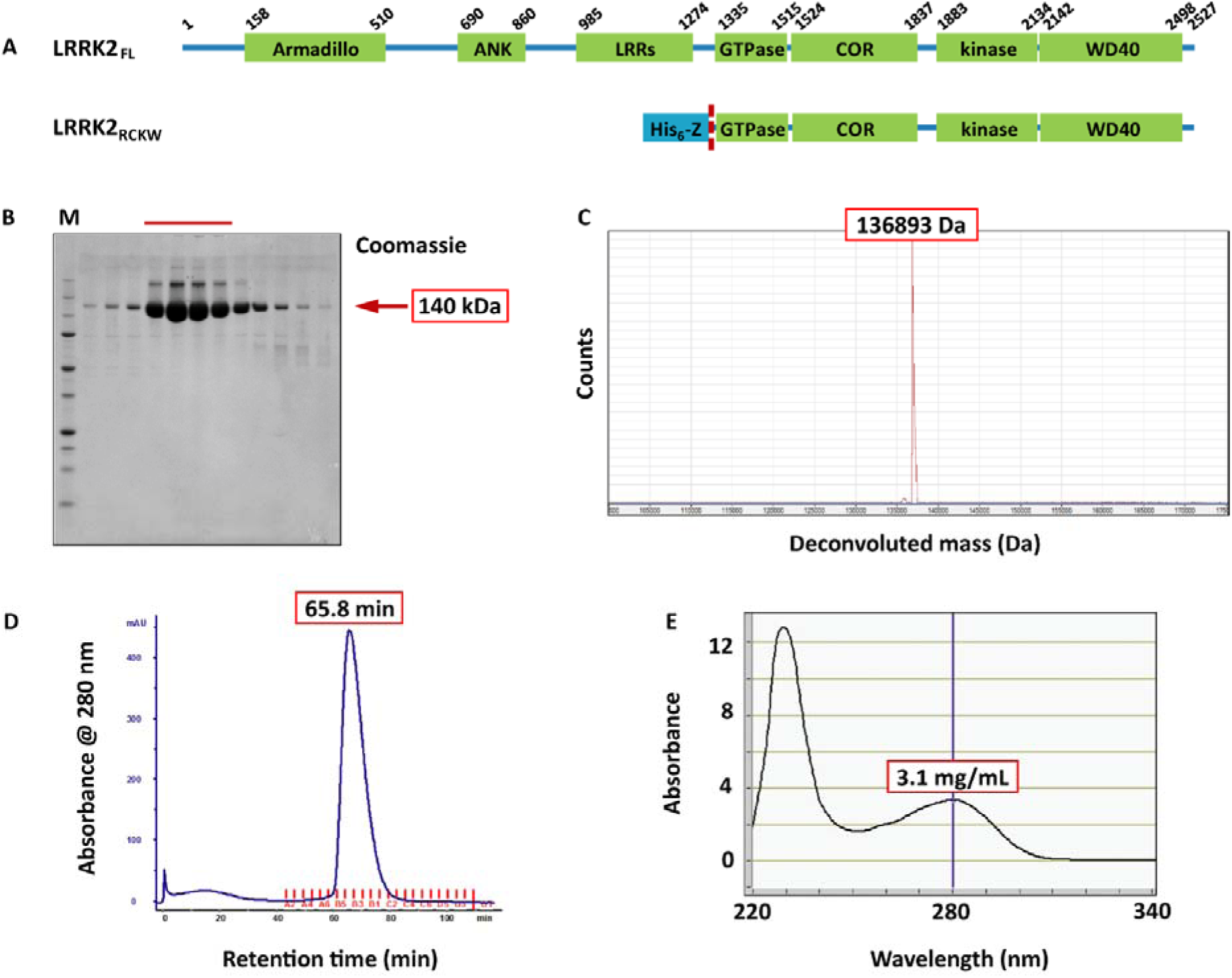
**A)** Domain architectures of the full-length LRRK2 protein and the His_6_-Z-tagged expression construct LRRK2_RCKW_. The dashed line indicates the TEV cleavage site for tag removal. **B)** LRRK2_RCKW_ was expressed in insect cells and purified to homogeneity. Shown is the SDS PAGE analysis of fractions from size-exclusion chromatography (SEC). SDS-denatured LRRK2 tended to migrate as dimers, trimers and tetramers in the gel. The red bar indicates the fractions that were combined and processed further. **C)** The purified protein was analysed by mass spectrometry. Shown is the deconvoluted spectrum. The only detected peak at 136893 Da corresponded well with the calculated LRRK2_RCKW_ mass of 136899 Da. Please note that peaks are generally broader and less accurate for larger proteins. **D)** The LRRK2_RCKW_ SEC retention time 65.8 min indicated dimer formation. The peak was reasonably defined, though slightly tailing, possibly due to a monomer dimer equilibrium. **E)** After SEC, the protein was concentrated to 3.1 mg/mL, as calculated from the absorbance at 280 nm. No signs of aggregation were detected at 340 nm.

### The most promising target for the treatment of Parkinson’s disease (PD)

LRRK2 is a genetic risk factor for PD, and ever since mutated LRRK2 was shown to cause the disease in 2002^*4*^, it has been speculated that the protein plays as well a role in the manifestation of the sporadic disease. This hypothesis is supported by several lines of evidence: First of all, carriers of LRRK2 mutations develop PD with late onset, slow progression and symptoms that are indistinguishable from non-carriers’ symptoms.^*5*^ Secondly, animal models confirmed that it is the triad of ageing, environmental factors and genetic predisposition that triggers PD, with every component being more or less influential, depending on the specific case.^*6*^ The effects of environmental factors depended on the cellular LRRK2 activity, with the highly active mutated LRRK2 leading to more pronounced effects in comparison to the non-mutated LRRK2.^*6*^ And as a proof of concept, membrane trafficking, the physiological process LRRK2 is a key regulator for^*2*^, ^*3*^, has been linked to several forms of neurodegeneration, including PD^*7*^. These findings suggest that modulating LRRK2 activity in PD patients might affect the progression of the disease in both carriers and non-carriers of LRRK2 mutations, which makes LRRK2 a worthwhile pharmaceutical target.^*8*^ At least some of the common mutations increase LRRK2’s kinase activity.^*9*^ In the past decades, the scientific community has developed an efficient toolkit for targeting kinases, notably in the field of cancer therapy. This will facilitate the development of LRRK2 kinase inhibitors as potential new drug candidates for PD.

### Towards an understanding of the LRRK2 molecule

To date no high-resolution structure of LRRK2 has been published. Even though significant progress has been made recently in describing the topology of LRRK2 using electron microscopy^*10*^, ^*11*^, the generated models have insufficient resolution for both rational inhibitor design and the understanding of the LRRK2 regulatory mechanism. We have established a robust method of expressing and crystallizing the most central portion of LRRK2 comprising the GTPase, the kinase as well as the WD40 domain (**Figure 1A**), but crystals diffracted poorly even after comprehensive optimization, limiting the diffraction to about 8 Å. Herein we describe our efforts to overcome this obstacle by crystallizing LRRK2 under microgravity conditions.

### Microgravity crystallization improves the diffraction properties of protein crystals

For determining the structure of a macromolecule by X-ray crystallography, it is absolutely crucial to work with the best possible crystal. The higher the order within the crystal, the higher the resolution of the final structure model. Diffraction data above 6 Å cannot be used for structure determination since such low-resolution data would result in uninterpretable electron density maps. The regularity of the macromolecule packing in a crystal is decreased by for instance a high solvent content, intrinsic flexibility of the crystallized macromolecule or by the incorporation of non-uniform particles into the crystal. Non-uniform particles derive of cause from impurities, but also from oligomers or aggregates, building blocks in a different conformation or orientation, and building blocks with decomposed or not bound co-factors. An even and long-lasting transport of building blocks to the site of crystal growth is maintained by diffusion. In contrast, convective currents and sedimentation disturb crystal growth and promote the incorporation of non-uniform particles.^*12*^ The only way of eliminating convective currents and sedimentation is to perform the crystallization experiment under microgravity conditions, as it is possible at the International Space Station (ISS). In the past decades, it has been shown for a multitude of crystallization systems, that microgravity improves the crystals’ diffraction properties.^*13*^ We therefore modified our crystallization procedure to allow for protein crystallization aboard the ISS. Advantages of the newly established method were that multiple crystallization conditions were tested simultaneously, that the reaction volumes were small, and that the integrity of the protein and crystallization solutions was maintained during transport because the samples were frozen. In addition, the astronaut was able to initiate the crystallization without special training, simply by moving the preassembled setup from a freezer to the temperature used for crystallization.

## Methods

### Cloning

The DNA coding for the LRRK2 residues 1327 to 2527 (taken from Mammalian Gene Collection) was PCR-amplified using the forward primer TACTTCCAATCCATGAAAAAGGCTGTGCCTTATAACCGA and the reverse primer TATCCACCTTTACTGTCACTCAACAGATGTTCGTCTCATTTTTTCA. The T4 polymerase-treated amplicon was inserted into the expression vector pFB-6HZB by ligation-independent cloning.^*14*^ According to Bac-to-Bac expression system protocols (Invitrogen), this plasmid was utilized for the generation of recombinant Baculoviruses.

### Expression and purification

Protein expression in Sf9 insect cells was performed as previously described.^*15*^ The protein construct contained an N-terminal His_6_-Z-tag, cleavable with TEV protease (**Figure 1A**). For LRRK2_RCKW_ purification, the pelleted Sf9 cells were washed with PBS, resuspended in lysis buffer (50 mM HEPES pH 7.4, 500 mM NaCl, 20 mM imidazole, 0.5 mM TCEP, 5% glycerol, 5 mM MgCl_2_, 20 μM GDP) and lysed by sonication. The lysate was cleared by centrifugation and loaded onto a Ni NTA column. After vigorous rinsing with lysis buffer the His_6_-Z-tagged protein was eluted in lysis buffer containing 300 mM imidazole. Immediately thereafter, the eluate was diluted with a buffer containing no NaCl, in order to reduce the NaCl-concentration to 250 mM and loaded onto an SP sepharose column. His_6_-Z-TEV-LRRK2_RCKW_ was eluted with a 250 mM to 2.5 M NaCl gradient and treated with TEV protease overnight to cleave the His_6_-Z-tag. Contaminating proteins, the cleaved tag, uncleaved protein and TEV protease were removed by another combined SP sepharose Ni NTA step. Finally, LRRK2_RCKW_ was concentrated and subjected to gel filtration in storage buffer (20 mM HEPES pH 7.4, 800 mM NaCl, 0.5 mM TCEP, 5% glycerol, 2.5 mM MgCl_2_, 20 μM GDP) using an AKTA Xpress system combined with an S200 gel filtration column. The elution time 65.8 min indicated the protein to be dimeric in solution. The final yield as calculated from UV absorbance was 1.2 mg LRRK2_RCKW_/L insect cell medium.

### Crystallization of the LRRK2_RCKW_/GDP/LRRK2-IN-1 complex

For sitting drop crystallization experiments, 20 μL precipitant solution each were added to the reservoirs of 3-well crystallization plates (Swissci). Altogether, approximately 600 different precipitant solutions were tested, covering a broad range of pH values, salts, additives and precipitants. With a Mosquito liquid handler (TPP Labtech), drops of a solution containing the protein-ligand complex (15 mg/mL LRRK2_RCKW_, 1 mM GDP, 500 μM LRRK2-IN-1) were transferred to the crystallisation plate, and mixed with aliquots of the respective precipitant solution. The volumes of protein complex solution and precipitant solution were 100+50, 75+75 and 50+100 nL, respectively. The plates were hermetically sealed and incubated at either 277 or 293 K. Crystal growth was documented using a Minstrel monitoring system (Rigaku).

### Microgravity crystallization within the US National laboratory aboard the International Space Station (ISS)

The microgravity crystallization experiments were carried out in the hardware Microlytic crystal former XL chip (Anatrace). For every experiment, 10 chips with 16 channels each were set up. Initially, the upper reservoirs including the diffusion channels were filled with 1.5 μL protein complex solution. Then the lower reservoirs were filled with 1.5 μL precipitant solution. After filling all 16 channels, the chip was sealed with adhesive foil and flash-frozen in liquid nitrogen. The protein complex solution was prepared from LRRK2_RCKW_ protein in storage buffer, with 1 mM GDP and 500 μM LRRK2-IN-1 added. The protein concentration was varied from 5 to 15 mg/mL. The precipitant solutions contained 0.1 M bis-tris-propane pH 8.0, 0.2 M potassium citrate and 10% ethylene glycol. In addition, the precipitant solutions contained 0 to 1.4 M NaCl and 10 to 30% PEG3350. The frozen chips were transported to ISS with the mission SpaceX CRS-12. To initiate crystallization, they were transferred to 293 K and incubated for 3 weeks. Following this, they were transferred to 277 K and transported back to earth for analysis.

### Crystal diffraction testing

The crystallization channels of the Microlytic chips were opened one by one with a cutter knife according to the manufacturer’s instructions. Immediately thereafter, a cryoprotectant solution was added to prevent the crystals from drying out. Individual crystals were then harvested with mounting loops (MiTeGen) and flash-frozen in liquid nitrogen. The diffraction properties of the frozen crystals were determined at Diamond Light Source (Oxford, UK) and at Advanced Light Source (Berkeley, CA), respectively.

## Results and discussion

### Making LRRK2_RCKW_ protein stocks

The expression construct LRRK2_RCKW_ was purified to homogeneity, as shown by SDS PAGE (**Figure 1B**) and mass spectrometry (**Figure 1C**). When subjected to size exclusion chromatography (SEC), it eluted in a single reasonably defined peak (**Figure 1D**), the retention time 65.8 min indicated the protein to form a dimer. All buffers used in the course of the purification were supplemented with GDP to stabilize the GTPase domain, the absence of a guanosine nucleotide led to LRRK2_RCKW_ precipitation even at protein concentrations below 1.0 mg/mL. The SEC elution fractions were pooled and concentrated to 3.1 mg/mL (**Figure 1E**).

### Preparing the LRRK2_RCKW_/GDP/LRRK2-IN-1 complex

Many kinases adopt an open, more flexible conformation in the absence of active site binders such as ATP or small-molecule inhibitors. This flexibility makes kinases unstable at higher concentrations and is generally disadvantageous for crystallization. A well-established approach is therefore to identify an active site binder such as a potent ATP competitive inhibitor first, and to crystallize the complex of the kinase and the stabilizing inhibitor. Since LRRK2 is regarded as a promising drug target^*8*^, a multitude of LRRK2 inhibitors have been developed^*16*^. One compound that in our hands strongly bound to the LRRK2 kinase domain, stabilized the protein and allowed for its crystallization was the small-molecule inhibitor LRRK2-IN-1^*17*^. It was added to the LRRK2_RCKW_ stock to a final concentration of 500 μM. At the same time, GDP was topped up to a final concentration of 1 mM. This mixture was concentrated to about 15 mg protein/mL. We assume the LRRK2_RCKW_ dimer still prevailed, with the GTPase domains saturated with GDP and the kinase domains saturated with LRRK2-IN-1. This complex was the starting point for our crystallization campaign.

### The LRRK2_RCKW_/GDP/LRRK2-IN-1 crystallization system

In initial sitting drop experiments several precipitant compositions turned out to allow for crystal formation. The morphologically most promising crystals were obtained with potassium citrate and PEG3350 at pH 8. By optimizing the concentrations of all components, we established a protocol for the reliable generation of LRRK2_RCKW_ crystals. The mainly lens-shaped crystals appeared overnight and did not change morphology after 5 days. They had a tendency of growing in layers, but single crystals with dimensions of 80x40 μm were also frequently obtained. The best crystals of each series were mounted in cryo loops, and their diffraction properties were determined at a synchrotron (Diamond Light Source, UK). We observed indisputable protein diffraction with, depending on the individual crystal, well-defined reflexions up to 8 Å (**Figure 2A**). This was a very promising result, taken into account that with a 4 or 6 Å dataset we will be able to see at least the domain arrangement in the LRRK2_RCKW_ dimer, which would help to understand the regulation of the key signalling molecule better. Indexing of the diffraction data with MOSFLM revealed that LRRK2_RCKW_ crystallized in space group P 2_1_ 2_1_ 2, the unit cell parameters were a=152, b=196, c=116 Å and α=β=γ=90°. From these parameters and the molecular weight of the protein, the asymmetric unit was likely to contain a LRRK2_RCKW_ dimer. The respective solvent content of 60% was rather high, but this corresponded well with the crystals’ fragility. Further attempts to improve the crystal quality such as streak seeding, hanging drop crystallization and additive screening were not successful. Yet another well-established, but logistically challenging way of improving the diffraction properties of a crystallization system is crystallizing under microgravity conditions.^*13*^ We considered it worthwhile in this case, because LRRK2_RCKW_ crystallization occurred reliably, and because diffraction just needed a subtle improvement (from 8 Å to below 6 Å), to allow for the collection of a valuable dataset.

**Figure 2.**
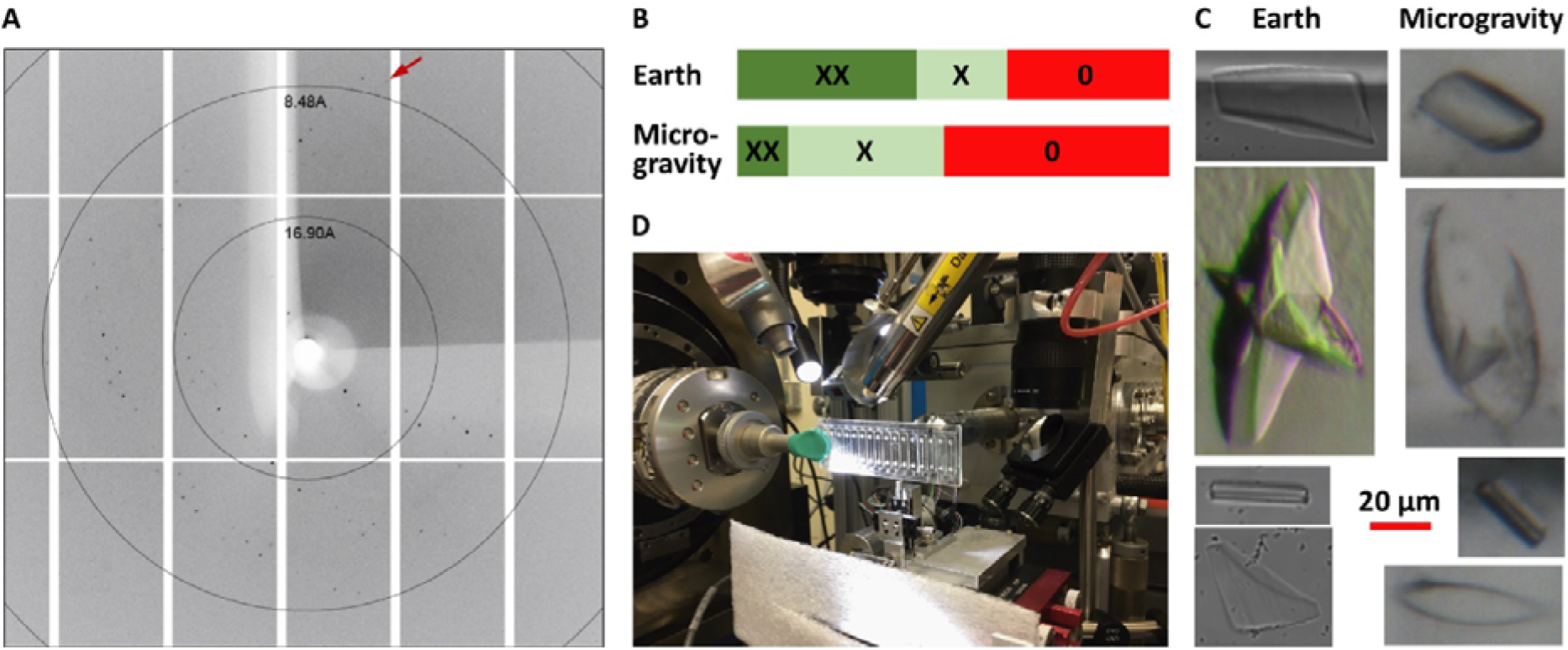
**A)**Diffraction pattern of earth-grown LRRK2_RCKW_ crystals. Diffraction is observed to at least 8 Å, as indicated by the red arrow. **B)** Both earth-incubated and microgravity-incubated crystal former chips contained crystals. The percentage of mountable XX crystals, however, was higher in the earth-incubated crystal former chips. X stands for small or layered crystals, and 0 stands for no crystals at all. **C)** The morphologies of earth-grown and microgravity-grown crystals were similar. Please note that differences in image quality derive from the microscopes used for documentation. **D)** A crystal former chip mounted to the goniometer at ALS beamline 5.0.3 (Berkeley) for room temperature diffraction testing.

### Growing LRRK2 crystals within the US National Laboratory aboard the International Space Station (ISS)

The 3-well sitting drop plates we had used for vapour diffusion crystallization were not suitable for crystallization under microgravity conditions. In the course of transporting the plates to the ISS and back, a certain amount of shaking is inevitable. This would lead to the precipitant solutions in the reservoirs merging with the crystallization drops, preventing crystal formation. Also, crystal growth would need to be delayed until the samples arrived at the ISS. We therefore performed the crystallization in the more robust Microlytic crystal former XL chips (Anatrace), with the nucleation being triggered by diffusive mixing in the embedded capillary-like channels. Another advantage of the crystal former chips was that they tolerated flash-freeze and thaw. This was tested several times, also at the Space Life Sciences Laboratory (Merritt Island) to demonstrate that our crystallization protocol reliably led to crystals of similar quality. For the microgravity experiment, the chips were set up, sealed and flash-frozen on earth, transported to ISS within the Dragon module on SpaceX-CRS-12 on August 14, 2017 while frozen at 178 K. After 3 days the experiment was transferred to the US National Laboratory for crystallization within the Merlin incubator at 293 K for 3 weeks before being transported back to earth aboard the SpaceX Dragon module. Two identical sets of chips were prepared, the only experimental difference was that one set was kept on earth and one set was sent to ISS, to explicitly examine the influence of microgravity on LRRK2_RCKW_ crystallization.

### Morphologies of earth-grown and microgravity-grown crystals

The experiment was carried out without incidents and without changes in protocol, and we recovered the undamaged ISS set of chips back for analysis on September 18, 2017 in Long Beach Airport. Initially, all 320 channels of both sets were inspected under the microscope and classified as either 0 (containing no crystals), X (low-quality crystals) or XX (mountable crystals, **Figure 2B**). Both conditions, the earth one and the microgravity one, yielded mountable crystals, highlighting the robustness of the experimental setup. Surprisingly, there were more channels with mountable crystals in the earth set than in the microgravity set (42% vs 12%). Another qualitative observation was that in the earth set, a lower initial precipitant concentration sufficed for crystal formation than in the microgravity set. An explanation for this might be that under microgravity conditions, the diffusive mixing of LRRK2_RCKW_ and the precipitant was much slower, so it took longer for a productive microenvironment to be established. And in the meantime, most of the protein was denatured and precipitated, thereby not suitable for crystallization anymore. While microgravity clearly affected the LRRK2_RCKW_ crystallization process, the morphology of the crystals, in terms of size and shape, was very similar in the earth set and the microgravity set (**Figure 2C**). There were huge lenses with irregularities in their centre (100 μm), slightly streaky shards with rough edges (50 μm) and tiny rods (20 μm). Hence, the goal to obtain bigger or morphologically more regular crystals was not achieved.

### Diffraction properties of microgravity-grown crystals

Next, we tested the diffraction properties of the crystals. 32 microgravity-grown crystals of diverse sizes and shapes were mounted in cryo loops and frozen for analysis at ALS beamline 8.2.1 (Berkeley). It turned out that none of these crystals diffracted to more than 10 Å. Therefore, no useful dataset could be collected. As a control, some of the Microlytic channels were left intact and tested directly at room temperature (ALS beamline 5.0.3, **Figure 2D**). The diffraction quality of the contained crystals was assessed by rastering with the beam along the channels. In this case, diffraction was limited to about 15 Å, indicating that it is not the freezing process that harms the crystals and decreases the diffraction.

### An explicit result – but no LRRK2 structure model

The generation of microgravity-grown LRRK2_RCKW_ crystals was successful in the first attempt, there were no technical problems in its course. The comparison of the microgravity-grown to the earth-grown crystals, however, clearly showed that microgravity conditions did not improve the diffraction properties of the crystals. Therefore, we failed to generate a LRRK2 structure model. There is a plethora of reports on how microgravity conditions improved protein crystallization.^*13*^ Why didn’t this approach help to produce better LRRK2_RCKW_ crystals? Protein crystallization is a multifactorial process, both the building block quality and the building block delivery define the intrinsic order of the resulting crystal. Microgravity mainly improves the latter, the building block delivery is even and long-lasting, convective currents and sedimentation are prevented. In contrast, the building block quality, in terms of uniformity and rigidity, is not affected by gravity. We assume that intrinsic flexibility of the protein, protein stability, multiple oligomerisation states and incorporation of non-uniform particles into the crystal prevented the formation of crystals good enough for the elucidation of the LRRK2_RCKW_ 3D model.

## Conclusions

LRRK2 is a multi-domain protein with a central role in the development of PD. Mechanistic understanding of the regulation of this complex protein and the rational design of selective LRRK2 inhibitors would be facilitated by high-resolution structures of full-length LRRK2 and its functional domains. Here we present data on the crystallization of the catalytically relevant domains of LRRK2 under normal and microgravity conditions. The crystallization method was successfully adapted to conditions that allow for transport to ISS and crystallization initiation with minimal manual effort. This was achieved by establishing a protocol where the crystallization setup was frozen in capillaries. This setup allowed storage at low temperature for several weeks and generated reproducible crystals with limited diffraction quality. Unfortunately, microgravity conditions did not improve diffraction quality of the crystals. The general experimental setup might however be useful for other projects aiming to crystallize proteins on board within the US National Laboratory of the ISS.

## Acknowledgements

SM and SK are grateful for funding support by the Michael J. Fox Foundation for Parkinson’s Research and the German cancer network DKTK. The Berkeley Center for Structural Biology is supported in part by the National Institutes of Health, National Institute of General Medical Sciences, and the Howard Hughes Medical Institute. The Advanced Light Source is a Department of Energy Office of Science User Facility under Contract No DE-AC02-05CH11231. In addition, the project was supported by the Center for the Advancement of Science in Space (CASIS) Award UA-2017–233, and we would like to thank Patrick O’Neill and Ken Shields of CASIS. We also thank Peggy Whitson of NASA and the team at SpaceX for providing the successful launch and return of our samples.

## Contributions

SM, JW, MA conducted experiments, MB, PR, AS, BF, SK supervised research, SM, SK wrote manuscript. All author edited the paper and approved the final version

